# Beyond the traditional simulation design for evaluating type 1 error rate: from ‘theoretical’ to ‘empirical’ null

**DOI:** 10.1101/311290

**Authors:** Ting Zhang, Lei Sun

## Abstract

When evaluating a newly developed statistical test, the first step is to check its type 1 error (T1E) control using simulations. This is often achieved by the standard simulation design S0 under the so-called ‘theoretical’ null of no association. In practice, whole-genome association analyses scan through a large number of genetic markers (*G*s) for the ones associated with an outcome of interest (*Y*), where *Y* comes from an unknown *alternative* while the majority of *G*s are *not* associated with *Y*, that is under the ‘empirical’ null. This reality can be better represented by two other simulation designs, where design S1.1 simulates *Y* from an alternative model based on *G* then evaluates its association with independently generated *G^new^*, while design S1.2 evaluates the association between permutated *Y^perm^* and *G.* More than a decade ago, Efron (2004) has noted the important distinction between the ‘theoretical’ and ‘empirical’ null in false discovery rate control. Using scale tests for variance heterogeneity and location tests of interaction effect as two examples, here we show that not all null simulation designs are equal. In examining the accuracy of a likelihood ratio test, while simulation design S0 shows the method has the correct T1E control, designs S1.1 and S1.2 suggest otherwise with empirical T1E values of 0.07 for the 0.05 nominal level. And the inflation becomes more severe at the tail and does not diminish as sample size increases. This is an important observation that calls for new practices for methods evaluation and interpretation of T1E control.

## 1 Introduction

Type 1 error (T1E) control evaluation using simulations is always the first step in understanding the performance of any newly developed statistical test. To formulate the problem more precisely, let us consider the current large-scale genome-wide association studies (GWAS) or next-generation sequencing (NGS) studies of complex and heritable traits. These studies scan through millions or more genetic markers (*G*s) across the genome for the ones associated with a trait of interest (*Y*), while accounting for environmental effects. Many *Y-G* association tests have been developed, and they often require the assumption of (approximately) normally distributed errors to maintain T1E accuracy, with some being more robust than others. For example, Bartlett test for variance heterogeneity has been shown to have large inflated T1E rates when the error term *e* follows a t-or *χ*^2^-distribution (Struchalin et al. 2010), and the likelihood ratio test (LRT) is similarly sensitive (Cao et al. 2014), while Levene’s test appears to be more robust (Soave et al. 2015; Soave and Sun 2017).

Standard T1E simulation design, denoted as S0, generates phenotype data *Y*_0_ ~ e under the ‘theoretical’ null model of no association, then independently generates genotype data *G* and estimates the empirical T1E rate from *Y*_0_ ~ *G* + *ϵ*; for notation simplicity and without loss of generality, intercept and additional covariates *Z*s are omitted from the conceptual expression of the regression model. A method is generally considered sound if T1E is well controlled under the *e* ~ *N*(0,*σ*^2^) assumption, and robustness is then evaluated by assuming other distribution forms for *e*. Given statistical accuracy of T1E control, statistical efficiency in terms of power will be studied by generating phenotype under an alternative, *Y*_1_ ~ *G* + *e*, and often it is assumed that *e* ~ *N*(0, *σ*^2^).

In practice, GWAS and NGS receive an *empirical Y* that comes from an unknown *alternative*, and a large number of *G*s of which the majority are *not* associated with *Y*. That is, most *Y-G* association pairs are in fact under the ‘empirical’ null. Now consider two alternative simulation designs to evaluate T1E control. Design S1.1 simulates *Y*_1_ ~ *G* + *e* from an alternative, then it independently generates *G^new^* and evaluates T1E from *Y*_1_ ~ *G^new^* + *ϵ*. Design S1.2 permutes the simulated *Y*_1_ and evaluates T1E from 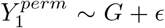. A important question can then be asked as to whether the S1.1 and S1.2 designs lead to similar T1E conclusion as the S0 design. In particular, even if the *e* ~ *N*(0, *σ*^2^) assumption was true and a test appeared to be accurate based on the S0 evaluation, do we expect it to perform well in real data which are better represented by the S1.1 and S1.2 simulation designs; note that *Y*_1_ is in contrast to *Y*_0_ and *ϵ* may or may be normally distributed. The answer would depend on the type of test statistics used.

Efron (2004) has brought up the discussion of the ‘theoretical’ vs. ‘empirical’ null more than a decade ago. Focusing on controlling the false discovery rate (FDR), Efron (2004) outlined several possible sources of non-normality including unobserved covariates and hidden correlation, and he proposed an empirical Bayes approach to the problem. Here, we study the practical implications of T1E evaluation based on the the commonly used ‘theoretical’ null simulation design S0 in the context of whole-genome scans. We show that while a method may appear to be accurate under S0 and assuming normality, it can have incorrect T1E rates under the ‘empirical’ null of S1.1 or S1.2 and also ‘assuming normality’. The fundamental cause of the discrenpancy is that, in evaluating 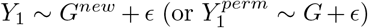, the marginal distribution of *Y*_1_ may not be normal even if it was generated assuming normality (*Y*_1_ ~ *G* + *e*), conditional on the true causal *G* and other covariates.

As a proof-of-principle, we will focus on scale tests for variance heterogeneity, recently proposed to identify *G*s associated with *variance* of a quantitative trait *Y* (Pare et al. 2010; Aschard et al. 2013; Cao et al. 2014; Soave et al. 2015). Traditional *Y-G* association tests focus on location parameters, studying changes in mean of *Y* across different genotype groups. Gene-environment (*G*x*E*) and gene-gene (*G*x*G*) are expected for complex traits. However, in practice, incomplete *E* data may preclude straightforward *G*x*E* interaction analyses, and computational or multiple hypothesis testing concerns can make whole-genome exhaustive *G*x*G* interaction searches undesirable. It was then recognized that because un-modelled interactions induce variance heterogeneity in *Y* when conditional only on *G*, scale tests such as Levene’s test, originally developed for model diagnostics, can be used to indirectly test for the interaction effects; it is worth noting that the causes of variance heterogeneity are multifaceted beyond potential interactions (Sun et al. 2013; Dudbridge and Fletcher 2014; Wood et al. 2014).

Inference of scale parameters is generally more sensitive than that of location parameters (Khan and Rayner, 2003). Thus, the distinction between the ‘theoretical’ and ‘empirical’ null can be particularly consequential for these emerging association tests that are designed to improve power by going beyond the first moment. In this work, we reveal the existing problems in T1E evaluation based on the ‘theoretical’ null simulation design S0. We show that (1) a T1E conclusion drawn from S0 could be different from the two alternative ‘empirical’ null simulation designs S1.1 and S1.2; (2) The T1E discrepancy can remain as sample size increases; (3) The T1E issue may be more severe at the tail.

In some settings, the ‘theoretical’ vs. ‘empirical’ null can also affect inference of location parameters in a regression, in addition to the better known cause of mean or variance model mis-specification. Assume *E* was available for direct modelling of the *G*x*E* interaction effect, Voorman et al. (2011) and Rao and Province (2016) showed that T1E rate of testing *G*x*E* or *G*x*G*_*non-repeating*_ in a whole-genome interaction scan can be sensitive to reasons beyond model mis-specifications. *G_non–repeating_* represents a fixed SNP *G* and we are testing its interaction with other SNPs, and *G*x*G*_*non-repeating*_ is statistically similar to *G*x*E* which we use, hereinafter, to refer to both. Focusing on inflated or deflated genomic inflation factor λ_*GC*_ (Devlin and Roeder 1999), Rao and Province (2016) demonstrated a larger variation in λ_*GC*_ (similar to a larger variation in T1E rates between different whole-genome association scans), when testing the interaction effect as compared to the main effect under the ‘theoretical’ null. They attributed this to dependence between the interaction test statistics, because *E* (or *G_no-repeating_*) is fixed between tests. And they noted that increasing sample size mitigates the problem. Here, we use this opportunity to revisit location testing of interaction effect. We show that, under the conventional ‘theoretical’ null, while T1E rates are indeed variable between simulation replicates, the average T1E rate is correct regardless of the sample size. In contrast, under the ‘empirical’ null, a different picture emerges as in the scale test setting above. In what follows, we first describe in Section 2 the scale tests to be investigated and the three simulation designs, S0, S1.1 and S1.2. We then provide numerical results from extensive simulation studies in Section 3, together with direct location tests for main and interaction effects. Importantly, we note that even if the departure from normality is generally minor in practice and appears to pass standard diagnostic tests for non-normality, the different null simulation designs can still noticeably affect conclusion regarding T1E control for some tests. Finally, in Section 4, we remark that future T1E evaluation and interpretation should go beyond the traditional ‘theoretical’ null and adopt the alternative ‘empirical’ null simulation designs.

## 2 Methods

For association study of a complex trait *Y* using a sample of size *n*, we first define genotype data *G_i_* for individual *i* at each SNP under the study. As in tradition, *G_i_* denotes the number of copies of the minor allele, coded additively as *G_i_* = 0, 1 and 2. And *G_i_* is assumed to come from a multinomial distribution, *G_i_* ~ multinomial(1, ((1 — *f*)^2^, 2*f*(1 — *f*), *f*^2^)), where *f* is minor allele frequency, MAF.

The analytical context of using scale tests to detect SNP *G* that influences variance of trait *Y* is the following. Suppose the true generating model is

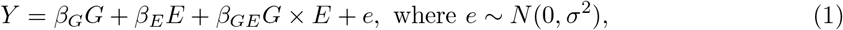

and suppose information regarding *E* was not collected, then the working model can only account for the main effect of *G*. However, it is straightforward to show that variances of *Y* stratified by the three genotype groups of *G* differ if *β_GE_* ≠ 0,

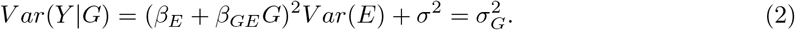

Thus, when *E* is missing and direct interaction modelling is not feasible, scale tests can be utilized to identify *G* associated with variance of *Y* (Pare et al. 2010). A joint location-scale testing framework can provide robustness against either *β_G_* = 0 or *β_GE_* = 0, and it can improve power if both main and interaction effects are present (Soave et al. 2015). Here we focus on studying the more sensitive scale tests, because the power of the joint test depends on the individual components.

Different scale tests have been studied in this context, and chief among them are the Levene’s test (Levene et al. 1960) considered by Pare et al. (2010) and Soave et al. (2015), and the LRT considered by Cao et al. (2014). Levene’s test for variance heterogeneity between *k* groups is an ANOVA of the absolute deviation of each observation *y_i_* from its group mean or median. The resulting test statistic Levene follows a F(*k* — 1, *n* — *k*) distribution under normality, and it is asymptotically 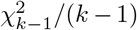 distributed; *k* = 3 in our case. Using median instead of mean to measure the spread within each group has been proved to be more robust to non-normality, particularly for t-distributed or skewed data (Brown et al. 1974; Soave and Sun 2017). And we will be using the median version of *Levene* in the remaining paper.

The variance likelihood ratio test considered by Cao et al. (2014) contrasts the null model of no variance difference with the alternative model,

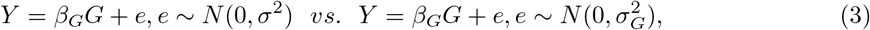

and conduct the corresponding LRT for 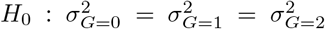. The corresponding test statistic *LRT_v_* is asymptotically 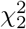, distributed; joint mean-variance LRT considering both *β_G_* = 0 and 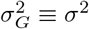 can be readily conducted. Cao et al. (2014) has pointed out that *LRT_v_* is sensitive to the normality assumption, but under normality they have demonstrated that *LRT_v_* has the correct T1E control. However, we show in the following that although this conclusion is analytically correct under the ‘theoretical’ null, it can be invalid when the method is applied to whole-genome scans which are better represented by the ‘empirical’ null.

Table 1 outlines the different null simulation designs, where S0 is the ‘theoretical’ null considered by Cao et al. (2014) as in convention, while S1.1 and S1.2 are the ‘empirical’ null designs that better represent the condition of real data. Under the S1.1 and S1.2 designs, the marginal distribution of phenotype *Y* is a weighted linear combination of normal distributions (Supplementary Materials). Thus, tests thought to be accurate based on S0 may have T1E issues based on S1.1 and S1.2, depending on the weighting factors and the means and variances of individual normal distributions. For example, *LRT_v_*, the LRT statistics for variance heterogeneity can be shown to be asymptotically equal to the weighted sum of independent 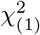 (Supplementary Materials and Theorem 3.4.1(1) of Yanagihara et al. 2005). Thus, before the simulation study in the next section, we shall expect that *LRT_v_* will have T1E issue when the simulated data is not normally distributed marginally.

**Table 1:**
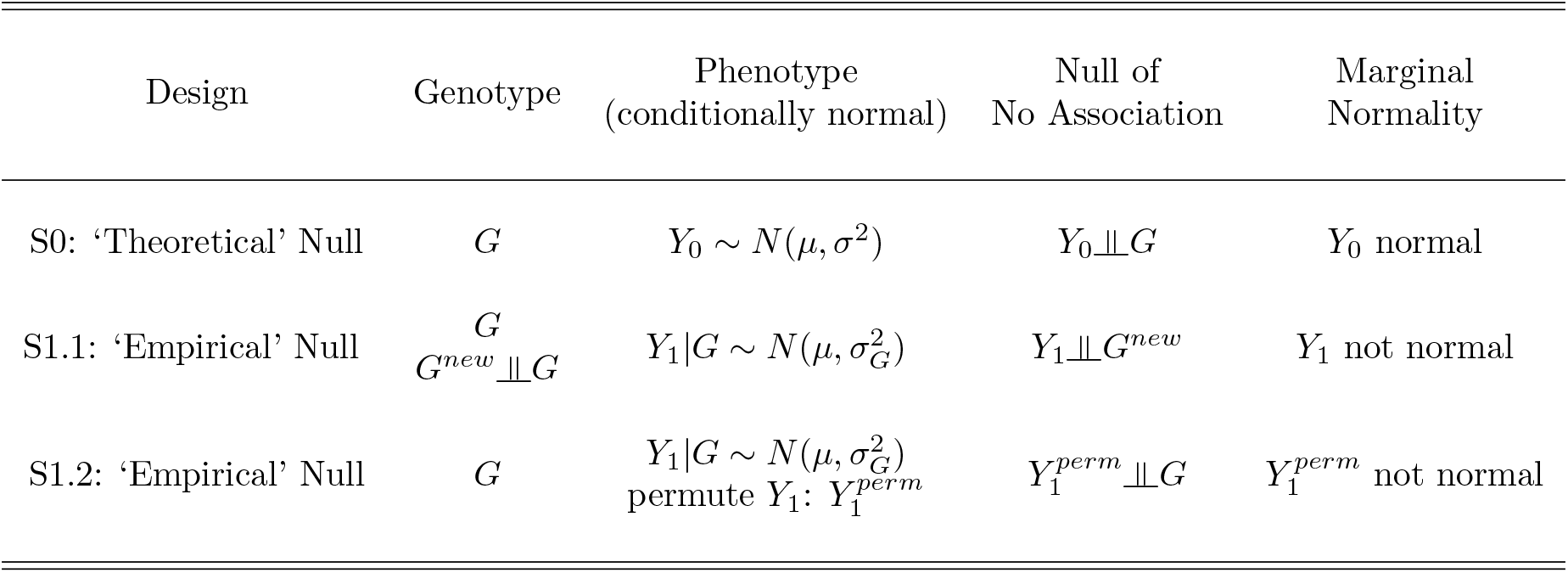
Summary of the three simulation designs, the ‘theoretical’ null S0, and the two ‘empirical’ null S1.1 and S1.2 for evaluating scale (or location) tests for *variance* (or mean) heterogeneity in phenotype *Y* across the three genotype *G* groups.

Assume that *E* was known, we can then directly test the interaction effect *β_GE_* using classical likelihood ratio test (*LRT_β_GE__*) or the score test (*Score_β_GE__*) based on model (1). In that case, it is straightforward to define the ‘empirical’ null design. That is, we first simulate *Y*_1_ = *β_G_G* + *β_E_E* + *β_GE_G* × *E* + *e* using model (1) based on the true *G* and *E*. We then independently simulate *G^new^* and test *β_GE_* from *Y*_1_ = *β_G_G^new^* + *β_E_E* + *β_GE_G^new^* × *E* + *ϵ*; similarly for S1.2.

There are a number of ‘theoretical’ null designs possible. For example, we can simulate *Y*_0_ = *β_E_E*+*e* without the main *G* effect (S0.1, Model I of Rao and Province 2016), or *Y*_0_ = *β_G_G*+*β_E_E* + *e* with the main effect (S0.2, Model II of Rao and Province 2016). In addition, we can also implement each ‘theoretical’ null model in two ways. Consider *Y*_0_ = *β_G_G* + *β_E_E* + *e*, we can simply simulate *nrep* sets of *G* and *E* to generate *Y*_0_. Alternatively, within each of *nrep.out* replicates (e.g. 100) of *E* in an outer simulation loop, we can simulate *nrep.in* replicates (e.g. 10^5^) of *G* and use them combined with the fixed *E* to simulate *nrep.in* replicates of *Y*_0_. We can then test *β_GE_* and estimate the T1E rate using the *nrep.in* replicates, similar to a whole-genome scan. Finally, we can average the T1E rate over *nrep.out* replicates to account for sampling variation inherent in simulation of *E* or one scan. We will be examining all four combinations for the ‘theoretical’ null. For the ‘empirical’ null, although the single-loop approach is possible, the *nrep.in* × *nrep.out* double-loop is more intuitive. That is, within each of *nrep.out* replicates of *G* and *E*, and *Y*_1_ simulated based on model (1), we simulate *nrep.in* replicates of *G^new^* for testing and T1E rate estimation. We then average across the *nrep.out* replicates.

## 3 Simulations

For evaluating scale tests for variance heterogeneity, we considered two modelling frameworks adopted, respectively, by Cao et al. (2014) and Aschard et al. (2013) (Tables 2). Cao et al. (2014) used model (3) to directly simulate variance heterogeneity in *Y* stratified by *G*. In contrast, Aschard et al. (2013) used model (1) to indirectly simulate variance heterogeneity that has better genetic epidemiology interpretation, because the size of *β_GE_* corresponds to power of scale tests under alternatives. Assume that *E* was known, model (1) also allows us to evaluate T1E control for our second study of directly testing for the interaction effect *β_GE_*. Conveniently, the corresponding ‘empirical’ null model S0 in Table 2, *Y*_0_ = *β_E_E* + *e*, is conceptually the same as the simulation model I of Rao and Province (2016), except *E* was *G_non–repeating_*.

**Table 2:**
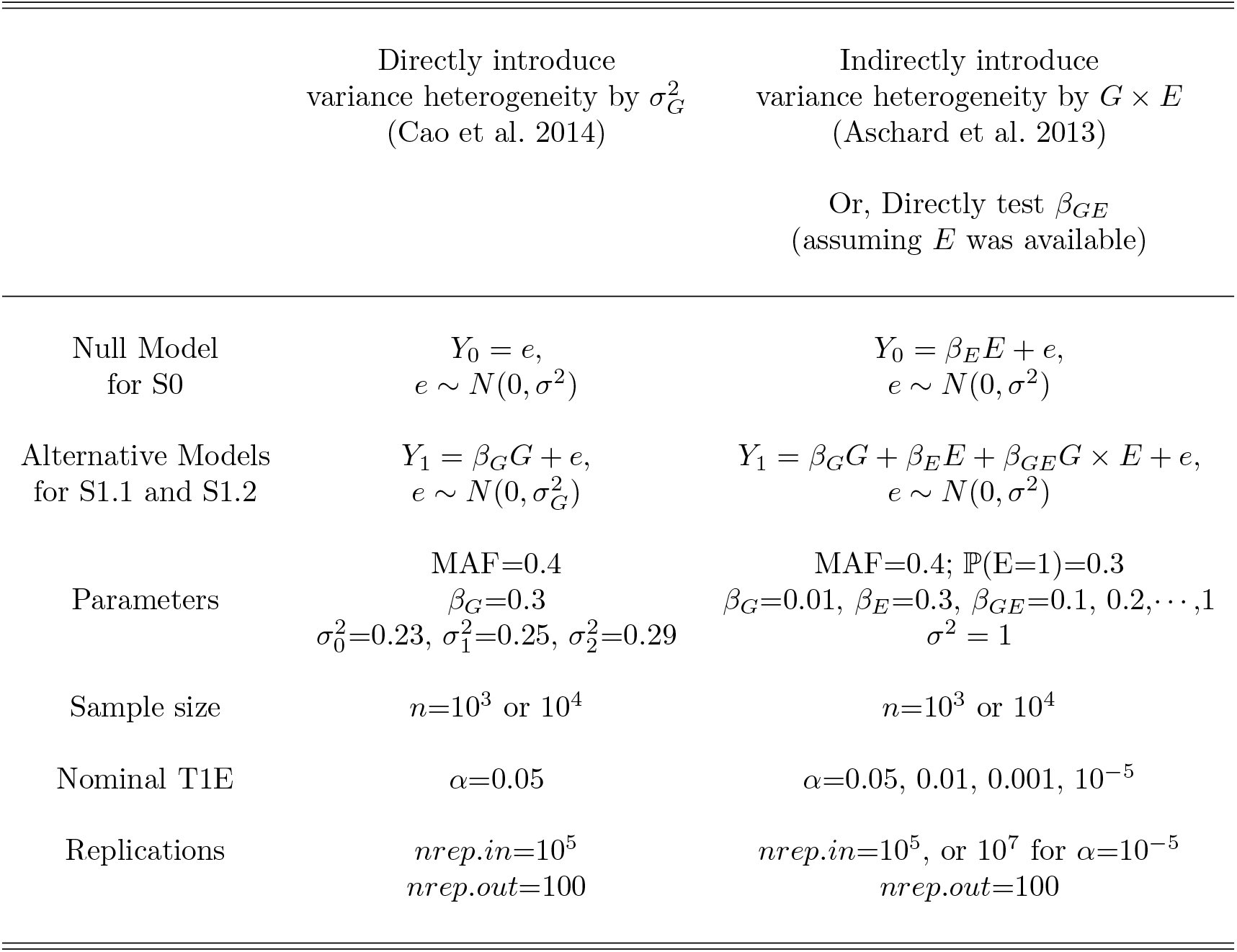
Summary of the two modelling approaches (Cao et al. 2014 and Aschard et al. 2013) used here to generate phenotypic variable *Y*_0_ under the ‘theoretical’ null simulation designs S0, and *Y*_1_ under the ‘empirical’ null simulation designs S1.1 and S1.2 as detailed in Table 1. Assume *E* was available for direct modelling and testing *β_GE_*, the Aschard et al. (2013) model coincides with Model I of Rao and Province (2016), except *E* was *G_non-repeating_*. T1E rate is first estimated from *nrep.in* simulation replicates in an inner loop (similar to one whole-genome scan), then averaged over *nrep.out* simulation replicates in an outer loop.

For each parameter value combination in Table 2, instead of studying power, we focused on evaluating T1E control of the LRT and Levene’s scale tests for variance heterogeneity, and location tests for interaction effect, by contrasting the proposed ‘empirical’ null with the previously considered ‘theoretical’ null. We first generated genotype and phenotype data for *G*, *G^new^*, (and *E* if needed), *Y*_0_, *Y*_1_ and 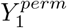 as described in Tables 1 and 2. We focus on the *nrep.in* × *nrep.out* double-loop implementation, but we note that the single loop design leads to the same conclusion as long as the total number of replicates is large (results not shown).

First, assume that information regarding *E* was not collected in practice, we applied the scale tests, *LRT_v_* and Levene, using the following working models,

- S0: *Y*_0_ ~ *G*
- S1.1, an alternative ‘empirical’ null: *Y*_1_ ~ *G^new^*
- S1.2, another alternative ‘empirical’ null: 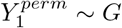

That is, we tested *Var*(*Y*_0_|*G*) across *G* under the ‘theoretical’ null of no association of S0, and *Var*(*Y*_1_|*G^new^*) across *G^new^* and 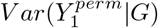 across *G* under the ‘empirical’ null of no association of, respectively, S1.1 and S1.2. We recorded the empirical T1E rates for each setting and bolded in red colour the ones that exceed the 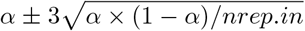 range, where *α* is the nominal T1E rate and *rep.in* is the number of simulation replicates used to estimate the empirical T1E rate for each of the *rep.out* replicates. Thus, 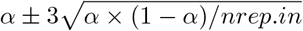 is a conservative interval. For completeness, we also kept the results of location tests (*LRT_m_* and *Score_m_*) for testing mean differences in *Y* across *G*, similarly contrasting the ‘theoretical’ null design of S0 with the alternative ‘empirical’ null designs of S1.1 and S1.2.

Revisiting the subtle dependency issue between interaction tests examined by Rao and Province (2016), we then assumed that *E* was available. That is, we tested *β_GE_* in *Y* = *β_G_G* + *β_E_E* + *β_GE_G* × *E* + *e* using the likelihood ratio test (*LRT_β_GE__*) and the score test (*Score_β_GE__*). However, S0 evaluated the association between *Y*_0_ and *G* × *E* under the conventional ‘theoretical’ null design, while S1.1 examined *Y*_1_ and *G^new^* × *E*, and S1.2 studied 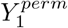 and *G* × *E^perm^* under the alternative ‘empirical’ null designs.

## 4 Results

As expected from the analytical insights, results in Table 3 show that while location tests for phenotypical mean differences (*LRT_m_* and *Score_m_*) are generally robust to the choice of ‘theoretical’ (S0) vs. ‘empirical’ (S1.1 or S1.2) null, it is not the case for the LRT scale test (*LRT_v_*) for variance heterogeneity; the empirical T1E rates of Levene’s test were slightly deflated but not significantly. Different choice of the null lead to different conclusions regarding the accuracy of *LRT_v_*. For example, simulation design S0 shows *LRT_v_* has the correct T1E control across the parameter values considered, but designs S1.1 and S1.2 suggest otherwise with empirical T1E values of 0.07 for the nominal α = 0.05 level for some settings. While the increased T1E rates under the S1.1 and S1.2 ‘empirical’ null designs appear to be mild and occur in extreme models (i.e. large un-modelled *β_GE_ G*x*E* interaction effect), results in Table 4 demonstrate that the T1E issue under the ‘empirical’ null simulation designs of S1.1 and S1.2 can be more severe at the tail. For example, for the nominal *α* = 1 × 10^−5^ level, the empirical T1E rate can be as high as 11.5 × 10^−5^. Because the genome-wide significance level for GWAS is *α* = 5 × 10^−8^ (Dudbridge and Gusnanto 2008), an inflation of false positive findings can be of a real problem in practice. Further, results in Table 5 confirm that increasing sample size *n* (from 10^3^ to 10^4^) does not mitigate the discrepancy in T1E conclusion drawn from the ‘theoretical’ vs. ‘empirical’ null. The root cause is that *Y*_1_ marginally is not normally distributed, even if it was generated (conditional on the true *G*) using a normally distributed error *e* term.

**Table 3:** Empirical T1E rates of *LRT_m_* and *Score_m_* location tests for mean difference in *Y* across the three *G* groups, and of *LRT*_*v*_ and Levene scale tests for variance difference in *Y*, based on the ‘theoretical’ null design of S0 and the alternative ‘empirical’ null designs of S1.1 and S1.2. Alternative empirical *Y*_1_ data were generated using the *Aschard*’*s genetic model* as described in Table 2. Empirical T1E rates outside 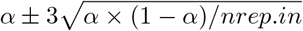 are bolded in red.

| $\alpha = 5 \times 10^{-2}, \quad n = 10^3$ | | | | | | | | |
| --- | --- | --- | --- | --- | --- | --- | --- | --- |
| | | $\beta_{GE}$ | 0.0 | 0.2 | 0.4 | 0.6 | 0.8 | 1 |
| Location | $LRT_m$ | S0 | 5.029 | 5.027 | 5.026 | 5.027 | 5.027 | 5.026 |
|  |  | S1.1 | 5.039 | 5.021 | 5.021 | 5.019 | 5.014 | 5.010 |
|  |  | S1.2 | 4.997 | 5.023 | 5.022 | 5.020 | 5.017 | 5.011 |
| | $Score_m$ | S0 | 5.002 | 4.998 | 4.997 | 4.998 | 4.998 | 4.997 |
|  |  | S1.1 | 5.014 | 4.992 | 4.993 | 4.991 | 4.985 | 4.981 |
|  |  | S1.2 | 4.974 | 4.994 | 4.993 | 4.990 | 4.988 | 4.983 |
| Scale | $LRT_v$ | S0 | 5.035 | 5.083 | 5.081 | 5.081 | 5.081 | 5.079 |
|  |  | S1.1 | 5.029 | 5.188 | **5.262** | **5.507** | **5.979** | **6.757** |
|  |  | S1.2 | 5.031 | 5.198 | **5.274** | **5.519** | **5.994** | **6.756** |
| | $Levene$ | S0 | 4.956 | 4.898 | 4.898 | 4.898 | 4.898 | 4.896 |
|  |  | S1.1 | 4.989 | 4.901 | 4.904 | 4.905 | 4.909 | 4.907 |
|  |  | S1.2 | 4.906 | 4.922 | 4.912 | 4.911 | 4.915 | 4.908 |

**Table 4:**
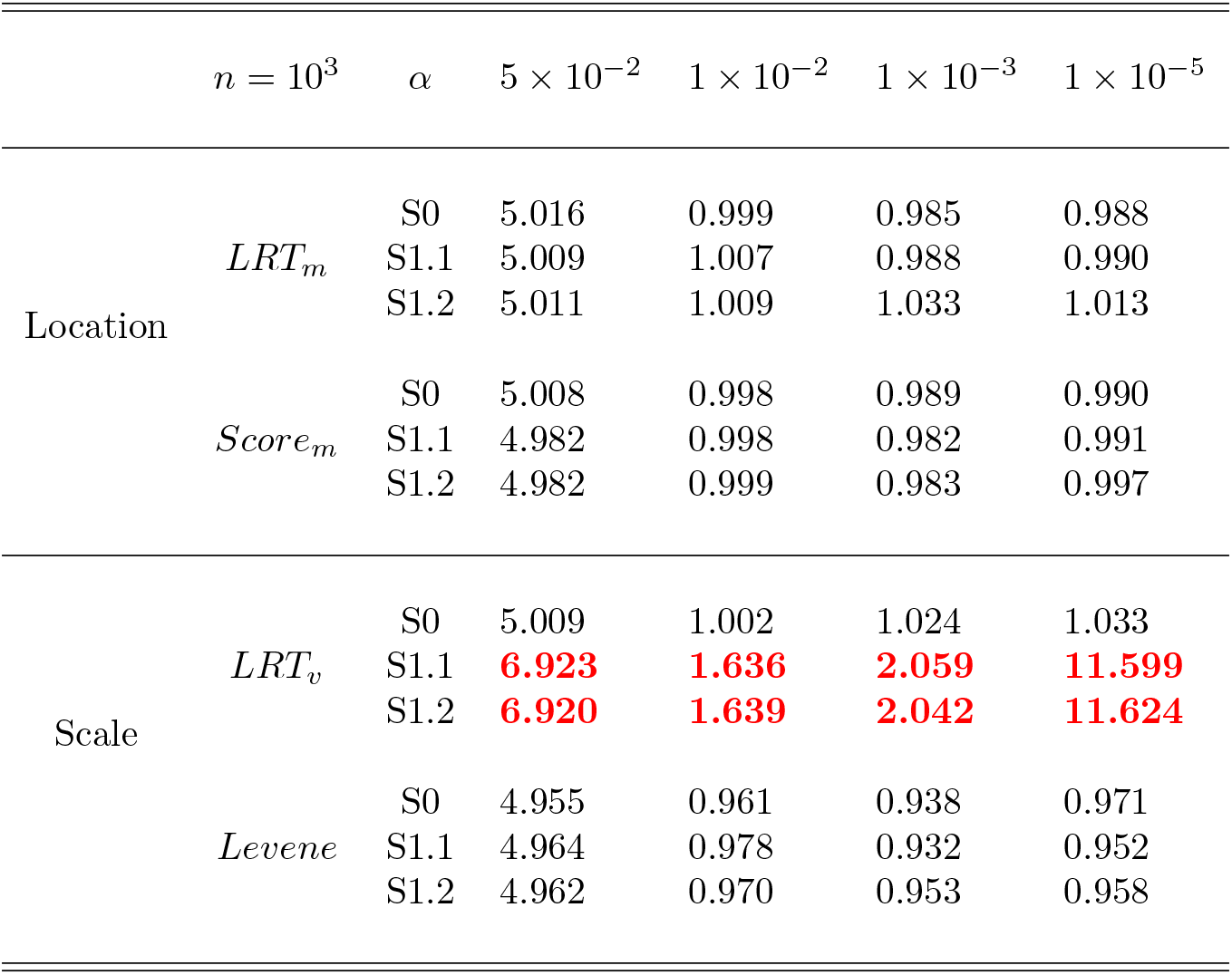
Empirical T1E rates of *LRT_m_* and *Score_m_* location tests for mean difference in *Y* across the three *G* groups, and of *LRT_v_* and Levene scale tests for variance difference in *Y*, based on the ‘theoretical’ null design of S0 and the alternative ‘empirical’ null designs of S1.1 and S1.2. Alternative empirical *Y*_1_ data were generated using the *Aschard’s genetic model* as described in Table 2, focusing on the extreme case of large interaction effect, *β_GE_* = 1. Empirical T1E rates outside 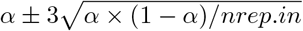 are boded in red.

**Table 5:** Empirical T1E rates of *LRT_m_* and *Score_m_* location tests for mean difference in *Y* across the three *G* groups, and of *LRT_v_* and *Levene* scale tests for variance difference in *Y*, based on the ‘theoretical’ null design of S0 and the alternative ‘empirical’ null designs of S1.1 and S1.2. Alternative empirical *Y*_1_ data were generated using the *Cao’s genetic model* as described in Table 2, and using two difference sample sizes of *n* = 10^3^ and 10^4^. Empirical T1E rates outside 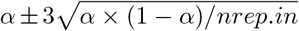 are bolded in red.

| $\alpha = 5 \times 10^{-2}$ | | | | |
| --- | --- | --- | --- | --- |
| | | | $n = 10^3$ | $n = 10^4$ |
| Location | $LRT_m$ | S0 | 5.011 | 5.012 |
|  |  | S1.1 | 4.992 | 5.003 |
|  |  | S1.2 | 4.934 | 4.989 |
| | $Score_m$ | S0 | 5.011 | 5.012 |
|  |  | S1.1 | 4.993 | 5.003 |
|  |  | S1.2 | 4.934 | 4.989 |
| Scale | $LRT_v$ | S0 | 5.103 | 5.165 |
|  |  | S1.1 | **7.034** | **7.125** |
|  |  | S1.2 | **7.007** | **7.020** |
| | $Levene$ | S0 | 4.924 | 4.945 |
|  |  | S1.1 | 4.905 | 4.965 |
|  |  | S1.2 | 4.874 | 4.825 |

In practice, it is routine (and recommended) to display and examine the empirical distribution of a trait under the study. However, Figure 1 shows that even under the most extreme setting where *β_GE_* = 1, the marginal histogram of *Y* appears to be approximately normal visually, unless a formal diagnostic test for normality was conducted. The slightly right-skewed empirical distribution of *Y* is the result of mixing six conditional distributions of *Y*, each perfectly normally distributed conditional on the causal *G* and *E*; this is the key difference between the ‘theoretical’ and ‘empirical’ null simulation designs, regardless of the sample size. For a less extreme case where *β_GE_* = 0.2, although both the histogram and Q-Q plot (Figure S1) suggest that normal distribution is a good fit (passing the Shapiro-Wilk normality test), the T1E discrepancy between the ‘theoretical’ and ‘empirical’ null remains albeit less severe as shown in Table 3 and Figure S2.

**Figure 1:**
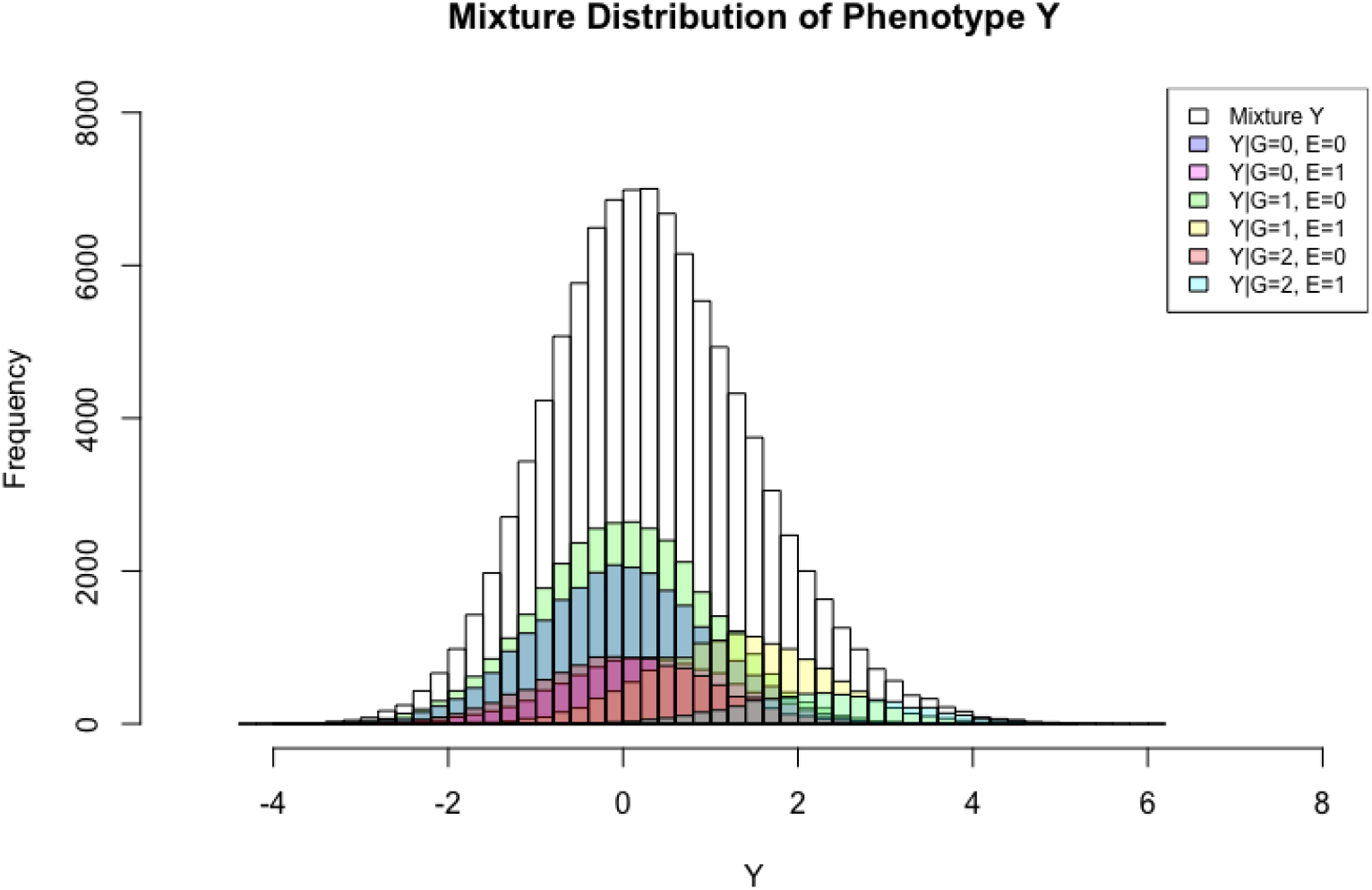
Marginal (mixture) and conditional (stratified by *G* and a binary *E*) histograms of empirical phenotype data *Y*_1_, based on the *Aschard’s model* as described in Table 2 when *β_GE_* = 1.

The asymptotic distribution of *LRT_v_* under the ‘empirical’ null is a weighted sum of 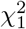 (Supplementary Materials). Figure 2 compares the asymptotic distribution (black solid curve) with the finite-sample distribution (red dashed curve) of *LRT_v_* under the ‘empirical’ null, as well as with 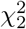 (blue dot-dashed curve), which is the asymptotic distribution of *LRT_v_* under the ‘theoretical’ null. While the asymptotic distribution derived under the ‘empirical’ null well approximates the finite-sample one, it is clear that the distributions of *LRT_v_* differ between the ‘empirical’ and ‘theoretical’ null; the difference is more visible on the scale of critical value for statistical significance (the vertical lines). Thus, applying *LRT_v_* to empirical GWAS or NGS while using the significance threshold derived from 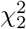 can lead to T1E problem.

**Figure 2:**
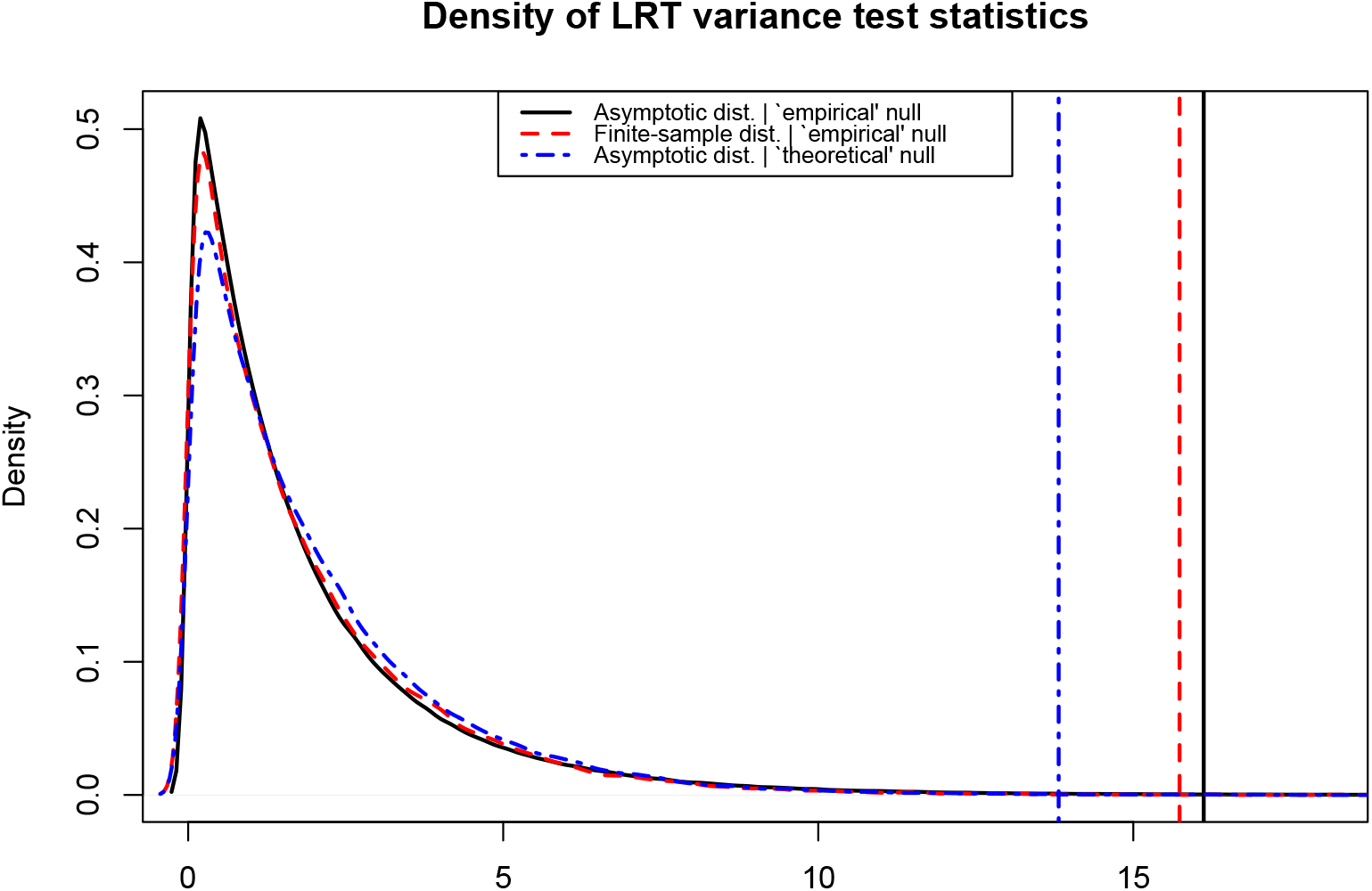
Comparison of the asymptotic distribution (black solid) and finite-sample distribution (red dashed) of *LRT_V_* under the ‘empirical’ null, with 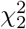 (blue dot-dashed) which is the asymptotic distribution of *LRT_v_* under the ‘theoretical’ null. Vertical lines correspond the 99.9% quantile cutoffs for *α* – 0.001.

Tables 3, 4 and 5 also included T1E results for testing phenotypic mean (as opposed to variance) difference across the genotype groups. Although location testing for the main effects are generally quite robust to the assumption of normality, problem can arise when testing for interaction effects beyond model mis-specification (Rao and Province 2016).

In testing the interaction effect *β_GE_* (*β_GGnon-repeating_* to be more precise), Rao and Province (2016) used the classical ‘theoretical’ null simulation design considering both S0.1 (without the main *G* effect) and S0.2 (with the main *G* effect). Regardless, Figures 1B-1C of Rao and Province (2016) showed that the variation in the resulting λ_*GC*_ was substantially bigger when testing *β_GE_* than testing *β_G_*. And their Figures 1D and 1E demonstrated that the variation diminishes as sample size increases. However, we note that this observation was made before averaging across the 414 simulated interaction scans/datasets; each scan contained 20,000 SNPs from which a λ_*GC*_ value was estimated.

The results of Rao and Province (2016) are consistent with ours shown in Figure S5. Figure S5 showed that scan-specific estimated T1E rates are indeed variable and become less so as sample size increases; 100 *G*x*E* interaction scans of 10^5^ SNPs each. However, it is important to note that the average T1E rate across *nrep.out* simulated scans reflects better the long-run behaviour of a method. Alternatively, assume *nrep.out* = 1, increasing the number of SNPs (i.e. the size of *nrep.in*) will decrease the sampling variation inherent in estimating T1E based on simulation studies. Indeed, results in Table 6 show that the T1E rate of testing *β_GE_*, estimated from 10^5^ × 100 (*nrep.in* × *nrep.out*) simulated replicates, is well controlled under the conventional ‘theoretical’ (S0) null simulation design. But, this is not the case for the ‘empirical’ (S1.1 or S1.2) null simulation designs. Similar to the *LRT_v_* scale test for variance heterogeneity, the discrepancy between two types of designs becomes more prominent at the tail and persists as sample increases (Table 6).

**Table 6:** Empirical T1E rates of *LRT_βGE_* and *Score_βGE_* location tests of the interaction coefficient *β_GE_ for the G* × *E interaction term in a regression*, based on the ‘theoretical’ null design of S0 and the alternative ‘empirical’ null designs of S1.1 and S1.2. Alternative empirical *Y*_1_ data were generated using the Aschard’s genetic model as described in Table 2 when *β_GE_* = 1, but *E* was assumed to be known in this case and direction interaction modelling was possible. Empirical T1E rates outside 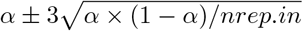 are bolded in red.

| | | $n = 10^3$ | | | $n = 10^4$ | | |
| --- | --- | --- | --- | --- | --- | --- | --- |
| $\alpha =$ | | $5 \times 10^{-2}$ | $1 \times 10^{-2}$ | $1 \times 10^{-3}$ | $5 \times 10^{-2}$ | $1 \times 10^{-2}$ | $1 \times 10^{-3}$ |
| $LRT(\beta_{GE})$ | S0 | 5.034 | 1.017 | 1.029 | 5.033 | 1.007 | 0.979 |
|  | S1.1 | **7.091** | **1.763** | **2.422** | **6.886** | **1.709** | **2.410** |
|  | S1.2 | **7.100** | **1.771** | **2.389** | **6.972** | **1.707** | **2.339** |
| $Score(\beta_{GE})$ | S0 | 4.982 | 1.003 | 1.002 | 5.021 | 1.004 | 0.972 |
|  | S1.1 | **7.028** | **1.738** | **2.363** | **6.874** | **1.705** | **2.400** |
|  | S1.2 | **7.040** | **1.747** | **2.339** | **6.960** | **1.703** | **2.326** |

## 5 Discussion

In this article, we highlight the importance of distinguishing the ‘theoretical’ and ‘empirical’ null distributions, first noted by Efron (2004), in a different application context. Focusing on scale tests for variance heterogeneity and through simulation studies, we showed that conclusions of type 1 error control of a statistical test could differ depending on the choice of the null. For example, the LRT variance test appears to be accurate under the ‘theoretical’ null but invalid under the ‘empirical’ null (Tables 3, 4 and 5, and Figure S2). Although the error term for generating the phenotype or outcome data was assumed to be normally distributed, the increased T1E rates under the ‘empirical’ null are, fundamentally, attributed to sensitivity of *LRT_v_* to departure from normality, because the marginal distribution of the empirical outcome data was not normal (Figures 1 and S1). Thus, tests shown to be sensitive to the assumption of normality are particularly vulnerable when applied to real data that are better represented by the ‘empirical’ null than the ‘theoretical’ null.

In practice, investigators often rely on visual inspection of histograms of outcome data as illustrated in Figures 1 and S1. And we have noted that the departure from normality does not have to be severe to have an effect on tests such as *LRT_v_*. For example, Soave et al. (2015) applied the *LRT_v_* test of Cao et al. (2014) to a GWAS of lung function measures in cystic fibrosis subjects. Despite the fact that the lung measures were approximately normally distributed and permuted prior to the variance association analysis, the histogram of GWAS p-values clearly showed an increased T1E rate (Figure S2.G of Soave et al. 2015); the actual application was a joint *LRT_m_* and *LRT_v_* test but the T1E issue was due to the *LRT_v_* component. Furthermore, for data appear to deviate from normal such as that in Figure 1, even if investigators chose to perform some standard normal transformations, the T1E issue can persist. For example, let us consider the phenotype data simulated based on Aschard’s genetic model, as described in Table 2 where *β_GE_* = 1 (Figure 1). After square-root or log transformations (Goh and Yap 2009), although the empirical marginal distribution of the phenotype improved as expected (Figure S3), the severity of T1E inflation of *LRT_v_* in fact worsened under the ‘empirical’ S1.1 and S1.2 null (Figure S4).

Beyond scale test of variance heterogeneity, Voorman et al. (2011) showed that spurious false positives can occur in genome-wide scans for *G*x*E* interactions, particularly in the presence of model mis-specification. And Rao and Province (2016) also presented inflated/deflated genomic inflation factors in a *G*x*G* interaction scan when one SNP is anchored (i.e. *G*x*G*_*non-repeating*_), using the conventional ‘theoretical’ null simulation design without any apparent model mis-specification. In our simulation studies, the situation when *E* was assumed available for direct modelling of the interaction term is similar to the dependency case examined previously. We note that the large variation in λ_*GC*_ estimate demonstrated by Rao and Province (2016) corresponds to the sampling variation inherent in estimating T1E rate from *nrep.in* replicates/SNPs across *nrep.out* replicates. This, however, does not translate to T1E issue based on the classical frequentist interpretation. Results in Table 6 show that, similar to scale test of variance, T1E conclusion for location test of interaction effect *β_GE_* is sensitive to the choice of ‘theoretical’ S0 vs. ‘empirical’ S1.1 or S1.2 null simulation designs. Theoretical justifications are provided in Section 3 of the Supplementary Materials.

In practice, permutation-based method must be carried out carefully, for example, in the presence of sample correlation (Abney 2015). Thus, the ‘empirical’ S1.1 design is perhaps earlier to implement than S1.2. For direct testing of the interaction effect *β_GE_*, the different ‘theoretical’ null designs (i.e. S0.1 without vs. S0.2 with main *G* effect) did not lead to different T1E conclusions.

To conclude, although we only presented two examples (i.e. scale tests for variance heterogeneity and location tests of interaction effects), the findings here have important implications for future evaluation of type 1 error control and interpretation. The newer test statistics being developed are increasingly complex, often going beyond the first moment such as the scale tests studied here, or beyond single variant approaches such as pathway and data integration analyses that have yet to be examined. Conventional simulation design S0 under the ‘theoretical’ null can lead to misleading conclusion regarding the accuracy of a test. The alternative simulation designs S1.1 and S1.2 under the ‘empirical’ null, on the other hand, can reveal the true behaviour of a test when applied to real data.

## Acknowledgements

The authors have no conflict of interest to declare. The authors would like to thank Dr. David Soave and Dr. Jerry Lawless for helpful discussions. This research was funded by the Natural Sciences and Engineering Research Council of Canada (NSERC, 250053-2013), and the Canadian Institutes of Health Research (CIHR, 201309MOP-310732-G-CEAA-117978) and to LS.

